# Cocaine-Enriched Oral *Streptococcus parasanguinis* Promotes Neuroimmune Dysfunction and Memory Impairment

**DOI:** 10.64898/2026.06.12.731966

**Authors:** Douglas Johnson, Tabinda Salman, Arshad Noorani, Barbara Benowitz, Yifei He, Kamala Sundararaj, Hannah Shelley, Zhenwu Luo, Zhuang Wan, Sylvia Fitting, Rachel D. Penrod, Wei Jiang

**Affiliations:** Department of Pharmacology and Immunology, Medical University of South Carolina, Charleston, USA, 29425; University of South Carolina, School of Medicine, Columbia, SC 29208; Department of Neuroscience, Medical University of South Carolina, Charleston, USA, 29425; Department of Psychology & Neuroscience, University of North Carolina at Chapel Hill, Chapel Hill, NC, USA, 27599; Ralph H. Johnson VA Medical Center, Charleston, USA 29403

**Keywords:** Cocaine use disorder, oral microbiome, oral-brain axis, bacterial metabolites, spatial memory, neuroinflammation

## Abstract

Chronic cocaine use is associated with neuroinflammation and cognitive dysfunction, but the underlying mechanisms remain unclear. We previously identified oral enrichment of *Streptococcus parasanguinis* (SP) and other species in individuals with cocaine use disorder (CUD), and here demonstrate that cocaine selectively enhanced SP growth *in vitro*. To investigate causality, antibiotic-pretreated wild-type C57BL/6 mice received chronic oral inoculation of SP, *S. salivarius*, *Neisseria flavescens*, or vehicle. SP-treated mice exhibited spatial memory impairment, increased brain IL-1β, and non-region-specific microglial activation, without detectable bacterial translocation into the brain. While amyloid-associated signaling changes were observed across all bacterial treatment groups, only SP induced cognitive deficits and neuroinflammation. Untargeted metabolomics identified distinct SP-associated oral-to-brain metabolite signatures, including cysteine S-sulfate (CSS) and altered histamine-associated metabolites. CSS and histamine induced neuroinflammatory and amyloid-associated responses *in vitro*. Together, these findings identify a cocaine-associated oral pathobiont that promotes neuroinflammation and neurodegeneration, suggesting a novel oral microbiome-brain axis in CUD.

## Introduction

Chronic cocaine use is associated with neuroinflammation and neurodegeneration, with smoked cocaine producing more severe and persistent impairments than intranasal use ^1–3^ and brain changes resembling Alzheimer’s disease (AD) ^4–7^. However, the underlying mechanisms remain unclear ^8–10^. These neuropathogeneses cannot be fully explained by cocaine’s direct effects. Notably, cocaine administration in humans reduces circulating IL-6 levels ^11^, while abstinence increases IL-6 ^12^, indicating a direct anti-inflammatory effect of cocaine. The conflicting reports of pro-versus anti-inflammatory effects of cocaine likely reflect differences in the exposure context ^11,13–18^ and suggest that chronic cocaine use-associated neuroinflammation may arise primarily through indirect mechanisms.

Previous studies from our team revealed that oral *Streptococcus* species are enriched in individuals with cocaine use disorder (CUD) who consume cocaine via smoking or snorting ^19^. In parallel, chronic cocaine use has been linked to impairments in attention, working memory, episodic memory, and executive function ^8,9,20^. These observations raise the hypothesis that cocaine smoking or snorting-associated oral microbial dysbiosis may influence neuroimmune and cognitive outcomes observed in chronic cocaine use.

In recent years, growing evidence has shown that the trillions of microbes inhabiting the gut and other mucosal surfaces are closely linked to brain health ^21,22^. A healthy microbiota produces a variety of neuroactive and immunomodulatory molecules that can shape mood, cognition, and behavior ^23–26^. These include a broad range of microbial-associated byproducts, such as microbiota-derived metabolites, outer membrane vesicles, and cell wall components such as lipopolysaccharides (LPS), lipoteichoic acid (LTA), and peptidoglycans (PGN) ^27^. Although the influence of the gut microbiome on mental health has been widely investigated, growing evidence, including our findings, highlights the importance of the oral microbiome-brain axis ^28–30^. The oral cavity harbors a complex microbial community, that can modulate systemic immunity and influence central nervous system (CNS) function through both direct and indirect mechanisms ^31–34^.

The oral cavity is closely linked to the brain through dynamic interactions among resident microbiota, systemic immune responses, and neural signaling pathways. As a result, the oral-brain axis has emerged as an important yet understudied contributor to neuropathogenesis, particularly in the context of smoking-or snorting-associated drug exposure. In this study, we tested the central hypothesis that the CUD-enriched oral microbiome identified in our previous work ^19^, particularly the salivary enrichment of *Streptococcus* species, promotes CNS immune activation and neuroinflammatory signaling, ultimately contributing to cognitive decline. To investigate the causal role of a cocaine-enriched oral microbiome in cognition and neuropathogenesis, wild-type C57BL/6 mice were orally inoculated and subsequently evaluated for neurobehaviors, neuropathology, and neuroimmune responses.

## Results

### Cocaine promotes the growth of *S. parasanguinis in vitro*

In our previous cross-sectional study profiling the salivary microbiome in CUD ^19^, *Streptococci* were the only genus enriched in the oral cavity (Fig. 1A), with the enriched species *S. parasanguinis*, *S. thermophilus,* and *S. australi* ^19^. Given the enrichment of *Streptococcus* species in the saliva of CUD, we next examined whether cocaine directly affects their growth. Notably, cocaine significantly enhanced the growth of *S. parasanguinis* (Fig. 1B). In contrast, the growth of *S. australis* and *S. thermophilus* remained unchanged (Suppl. Fig. 1B). No difference was observed between the two cocaine concentrations tested (Suppl. Fig. 1A). The lower concentration was selected because it more closely reflects the physiological levels detected in cocaine users ^35^. These findings indicate that cocaine selectively promotes the growth of *S. parasanguinis*, potentially explaining its enrichment in the oral microbiome of CUD.

**Figure 1.**
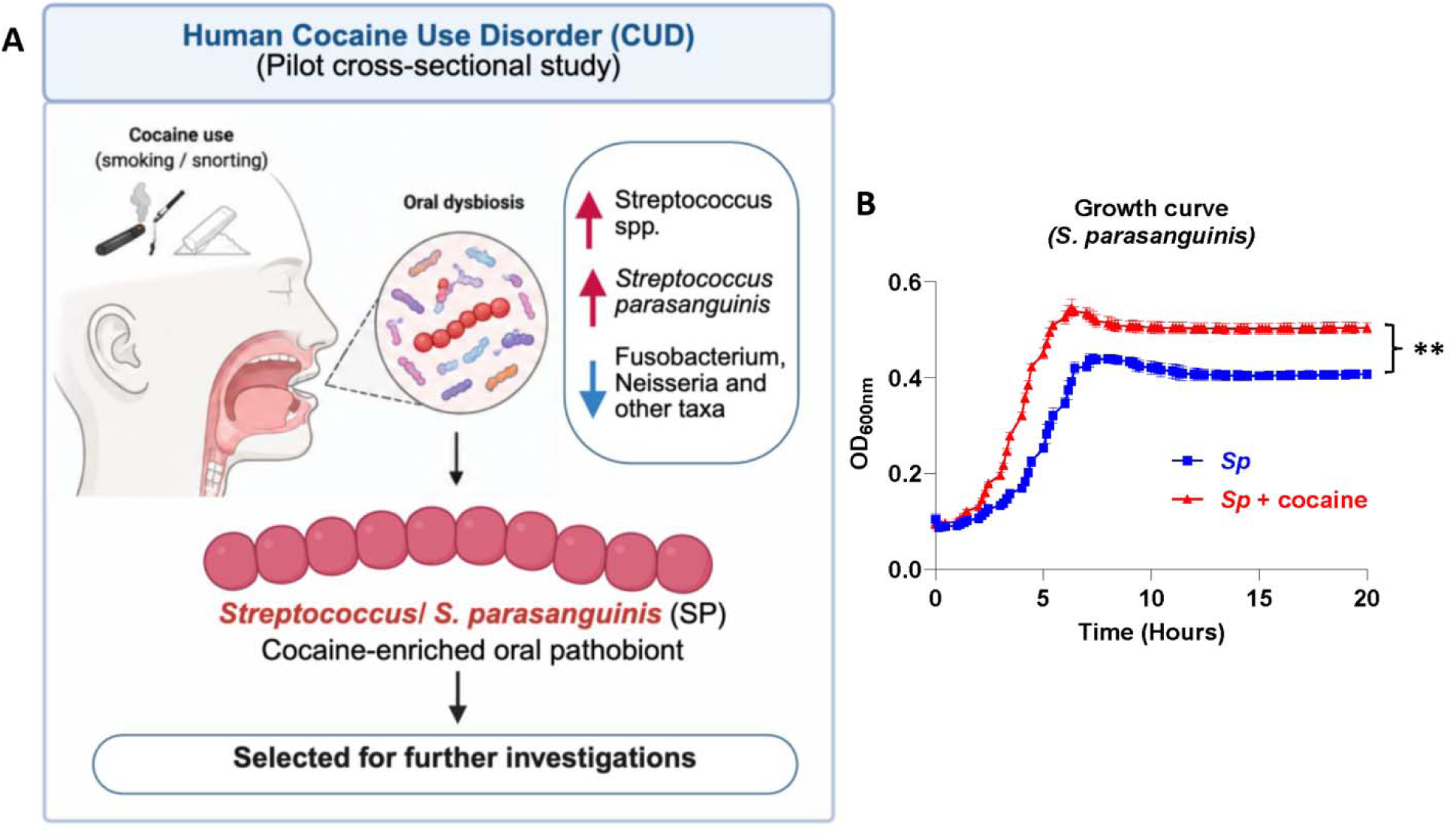
Cocaine promotes the growth of *S. parasanguinis in vitro.* **(A)** A schematic of our preliminary finding shows enrichment of *Streptococcus parasanguinis (*SP) in the oral cavity of chronic cocaine users, along with a few other Streptococcus species ^19^. **(B)** SP, along with the other three *Streptococcus* species, was cultured with cocaine at 5 µg/mL in vitro, supplemented with glucose (50 mM) in ZMB1 medium. The bacterial growth curve was assessed using optical density (OD600), measured in triplicate. SP exclusively showed an enhanced growth pattern in the presence of cocaine but not the other *Strep.* species (*supplementary* fig. 1).

### Oral *S. parasanguinis* inoculation leads to spatial memory decline

To investigate the causality of cocaine-enriched *S. parasanguinis* and oral-to-brain mechanisms, C57BL/6 mice were pretreated with mixed antibiotics first, and then orally inoculated for 3 months with *S. parasanguinis* (SP, enriched in the saliva of cocaine smokers), *S. salivarius* (SS, *Streptococcus* species control), and *Neisseria flavescens* (NF, diminished in the saliva of CUD), alongside a vehicle control group receiving carboxymethylcellulose (CMC). This experimental design was intended to model the sustained microbial shifts observed in CUD (Fig. 2A). To confirm bacterial colonization, oral isolates recovered at the study endpoint were characterized using microbiological and biochemical analyses, which verified successful oral colonization following repeated inoculations (Suppl. Fig. 2). Following confirmation of successful colonization, body weight and serum corticosterone levels were monitored to rule out potential effects of environmental stress (Suppl. Fig. 2). Mice were then subjected to a battery of behavioral assessments evaluating recognition memory, anxiety-like behavior, and depression-like behavior, none of which showed significant differences among groups (Suppl. Fig. 3), except for the Barnes maze test assessing spatial learning and memory. In the Barnes maze task, mice underwent training to locate their escape chamber over five consecutive days. During the test, SP-treated mice spent more time in the escape hole zone (Fig. 2D) and the escape quadrant (Fig. 2E) compared with vehicle controls. Additionally, SP-treated mice exhibited more entries into the target area (Fig. 2G) and a longer latency to reach the escape chamber (Fig. 2C), indicating that they required more trials to locate the escape chamber in the Barnes maze. Heatmaps and representative tracking plots further supported these behavioral divergences, showing that SP-treated mice exhibited broader search trajectories and increased occupancy outside the target quadrant during the test trial (Fig. 2H and 1I). Together, these findings suggest that prolonged oral exposure to SP selectively impaired spatial search strategies and memory performance.

**Figure 2.**
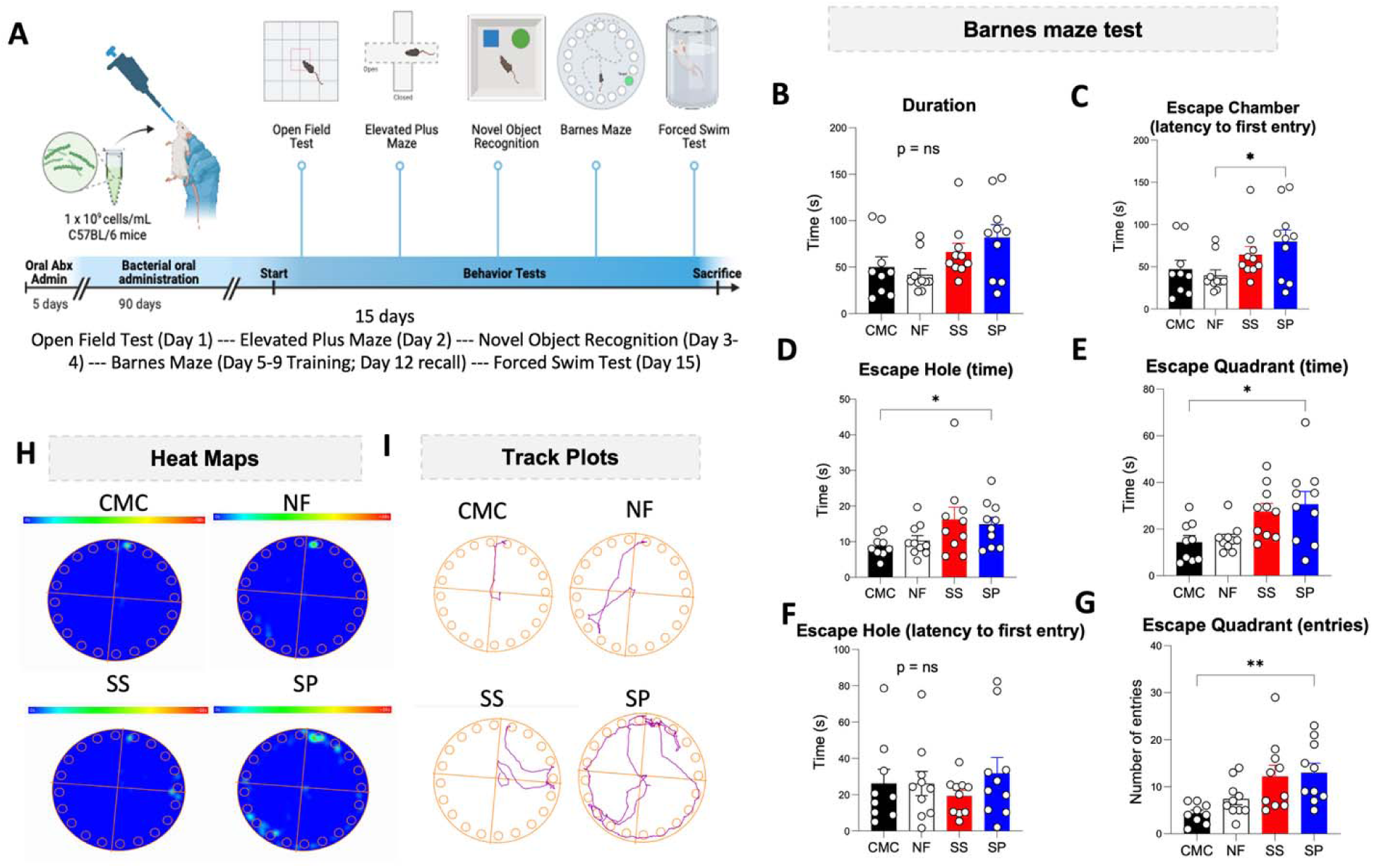
Chronic oral administration of *S. parasanguinis* impairs spatial memory performance in the Barnes Maze. **(A)** Schematic of the mouse study design. C57BL/6 mice received oral antibiotic pretreatment followed by repeated oral administration of *Streptococcus parasanguinis* (SP), *Streptococcus salivarius* (SS), *Neisseria flavescens* (NF), or vehicle control (CMC). Behavioral testing was conducted after completion of bacterial administration, followed by sacrifice and tissue collection. Various parameters in the Barnes-Maze test, indicating the memory decline in the SP-treated mice, i.e., **(B)** total trial duration (F*_(3,35)_* = 1.303, p =0.2888), **(C)** increased latency to first entry into the escape chamber (F*_(3,35)_*= 1.285, p =0.043), **(D)** increased time spent investigating the escape hole (F*_(3,35)_* = 1.896, p= 0.0465), **(E)** elevated time spent in the escape quadrant (F*_(3,35)_* = 2.843, p= 0.0057), **(F)** latency to first entry in escape hole (F*_(3,35)_*= 0.7829, p =0.6301), and **(G)** more number of entries into the escape quadrant (F*_(3,35)_* = 3.728, p =0.0036) in SP-treated mice as compared to control group. **(H-I)** Representative trajectories and heatmaps from the Barnes maze on test day, demonstrating the spatial search strategies across treatment groups. Data are presented as mean ± SEM (n = 9-10 per group). *p < 0.05, **p < 0.01; one-way ANOVA with Tukey’s multiple comparisons test.

### Oral *S. parasanguinis* inoculation induces non-region-specific microglia activation and neuroinflammation without evidence of systemic macrophage infiltration to the CNS

Previous studies have shown that peripheral macrophages can infiltrate the CNS following injury, infection, or other conditions that disrupt the blood-brain barrier ^36^. We next investigated whether chronic oral inoculation with SP promotes infiltration of peripheral macrophages into the CNS and influences CNS myeloid cell activation. Using fluorescent RNA in situ hybridization (ISH) on sagittal FFPE mouse brain sections, we assessed IBA1 expression as a marker of microglia activation. SP-treated mice exhibited significantly increased IBA1 expression compared with vehicle controls, indicating robust microglia activation in a non-region-specific manner, while SS-treated mice showed a similar but less pronounced increase (Fig. 3A and B). Notably, elevated IBA1 signals were broadly distributed throughout the brain rather than restricted to a specific anatomical region following SP inoculation. TMEM119 was analyzed as a marker to distinguish macrophages (negative) from resident microglia (positive) ^37^. Careful examination of the brain sections revealed no prominent TMEM119-negative/IBA1-positive cell populations. Instead, all IBA1+ cells co-localized with TMEM119 (Fig. 3C), suggesting that the observed increase in IBA1 signal primarily reflected activation of resident microglia rather than infiltration of peripheral macrophages.

**Figure 3.**
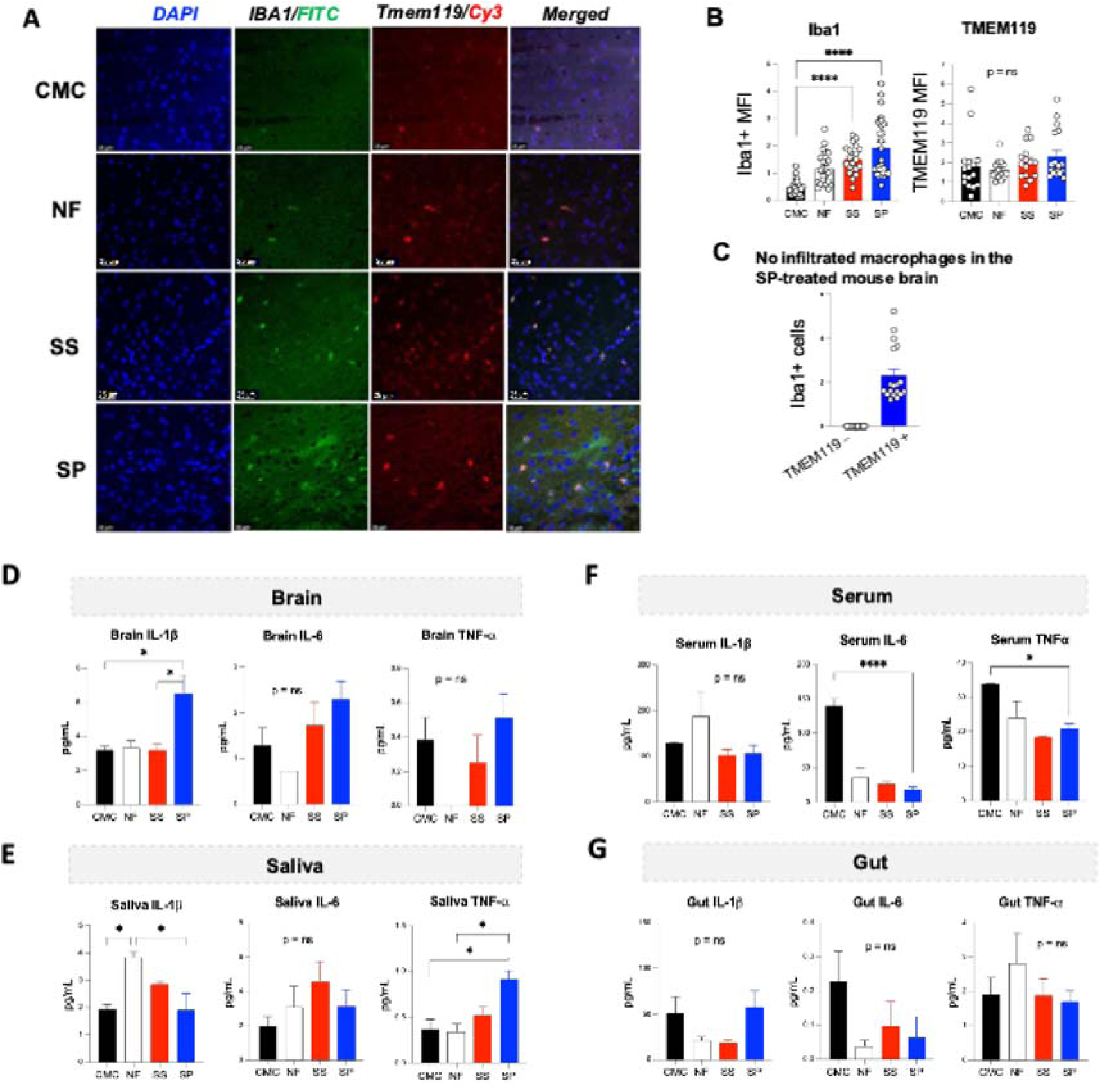
Chronic oral inoculation of *S. parasanguinis* triggers microglial activation and induces neuroinflammation without systemic macrophage infiltration in the brain. (A) RNA-based *in situ* hybridization (RNAScope). Representative images of 5 μm thick sections from the mouse sagittal brain embedded in FFPE. The scale bar is 25μm in the cortex of the brain. Sections were stained with IBA1 (activated microglia/macrophages, FITC) and TMEM119 (Red channel Cy3); a distinguishing marker for macrophages (negative) to microglia (positive), and DAPI as a counterstain. **(B-C)** Quantification of the mean fluorescent intensities (MFI) through ImageJ/Fiji. Microglia (Iba1 and TMEM119) activation was observed/increased in the *S. parasanguinis* inoculated group as compared to the control groups. Also, the MFIs were compared for IBA1 versus TMEM119. No prominent cluster of Iba1+ cells was detected that was not co-localized with TMEM119+, showing activation of microglia but not infiltration of systemic macrophages into the brain. **(D-G)** Protein levels of pro-inflammatory cytokines measured by MSD multiplex immunoassay in mouse brain lysates, saliva, serum, and gut lysates. Analytes include IL-6, IL-1β, and TNF-α as indicated in each panel. No significant increase was observed between the *S. parasanguinis*-treated group and vehicle controls in the serum and gut samples, while the brain IL-1β and saliva TNF-α levels were found to be increased in the SP treatment group compared to the control. n = 3-5 animals per group; sample size varied due to sample availability, and all available samples were used; *p < 0.05, **p < 0.01, ***p < 0.001, ****p < 0.0001; one-way ANOVA, followed by Tukey’s multiple comparison test.

To further evaluate proinflammatory responses across tissue compartments, three typical proinflammatory cytokine protein levels were measured in brain, saliva, serum, and gut samples by multiplex immunoassay. In the brain, SP-treated mice exhibited significantly elevated Il-1β levels relative to controls, whereas IL-6 and TNF-α showed a trending increase (Fig. 3D). In the oral cavity, IL-1β was highest in the NF-treated mice versus SP and CMC, TNF-α was significantly increased in SP-treated mice compared with controls, while salivary IL-6 was similar among the four study groups (Fig. 3E). In serum, decreased IL-6 and TNF-α levels were observed in SP-treated mice versus controls with no changes in IL-1β (Fig. 3F). Likewise, proinflammatory cytokine levels in the gut were similar among the four study groups (Fig. 3G). These results indicate that prolonged oral exposure to SP is associated with neuroinflammation, whereas no evidence of broad peripheral and intestinal inflammation was observed. This is notable given that gut inflammation can influence enteric and gut-brain signaling pathways, which might otherwise confound behavioral interpretation ^38^. Collectively, these findings identify a proinflammatory signature in the brain strictly associated with chronic oral SP exposure, characterized by increased microglial activation, without corresponding evidence of broad systemic or gut inflammation.

### Oral bacterial exposure perturbs APP-processing pathways and increases brain amyloid-**β** (A**β**) levels in all three microbiome groups compared to CMC

Given the neuroinflammation and cognitive impairment observed following oral SP exposure, we next examined whether SP administration affected molecular pathways related to the Amyloid Precursor Protein (APP) processing pathway and Aβ production in the brain (Fig. 4A). qPCR analysis of whole-brain lysates revealed reduced expression of *Psen1* (PSEN1 protein is the catalytic core of the γ-secretase complex ^39^) and *Bace1* (BACE1 protein is the first cleavage of APP ^39^) in SP-treated mice relative to CMC controls. In contrast, APP and *Tau* expression remained unchanged across groups (Fig. 4B). In contrast, *Gsap* expression, a protein promoting amyloidogenic APP processing ^40^, was increased in SS-and slightly increased in SP-treated mice compared with controls (Fig. 4B). These findings suggest that chronic oral bacterial exposure alters transcriptional components of amyloid-related processing pathways in the brain.

**Figure 4.**
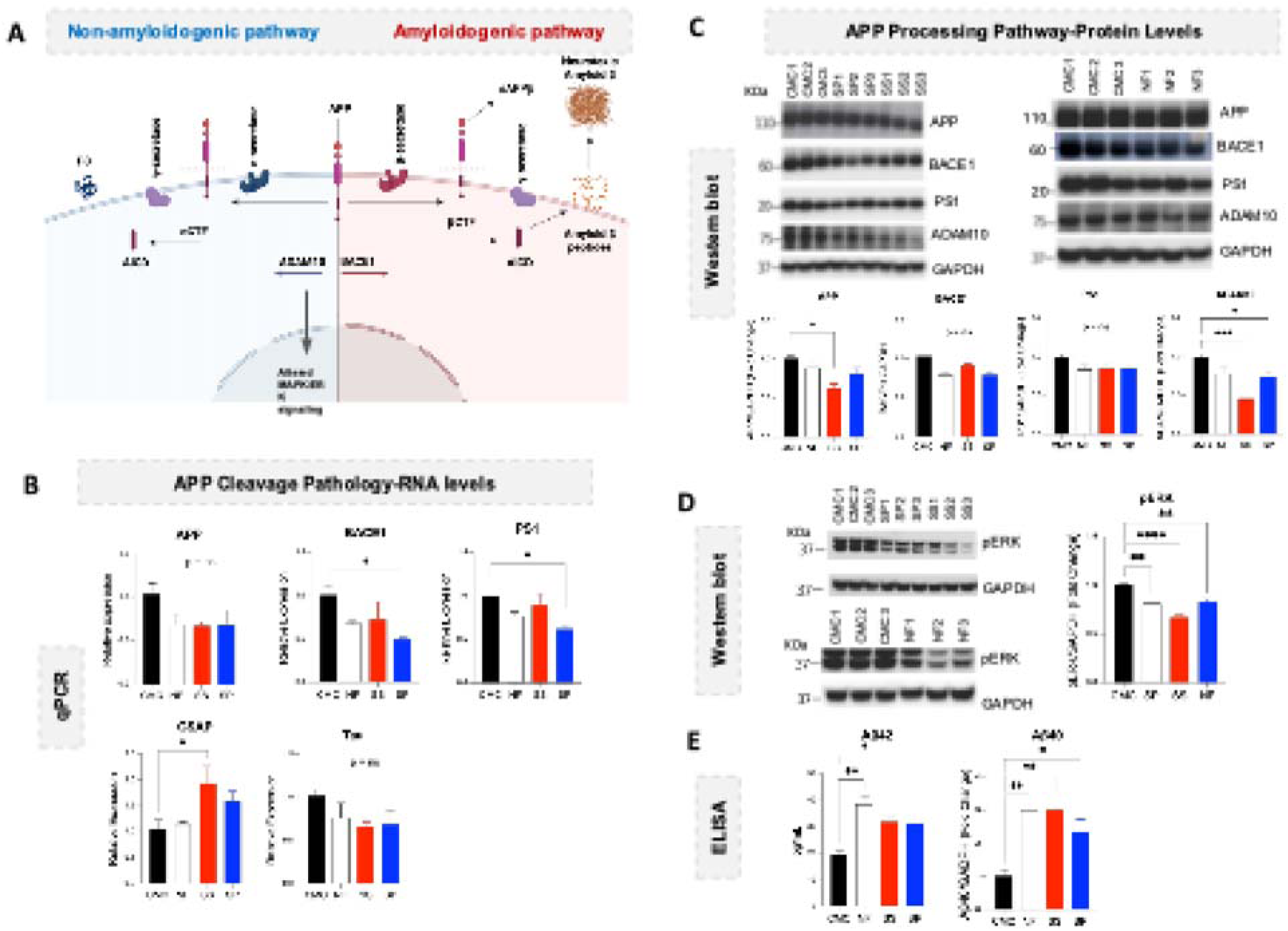
Modulation of amyloidogenic processing and amyloid-β production following chronic oral bacterial administration versus CMC controls, but not specific to *S. parasanguinis*. (A) Schematic of APP processing pathways showing two pathways. In the non-amyloidogenic pathway, APP is cleaved by α-secretase (ADAM10); in the amyloidogenic pathway, sequential cleavage by β-secretase (BACE1) and γ-secretase (PS1) generates amyloid-β peptides. **(B)** qPCR analysis of amyloid-related transcripts in whole-brain lysates. Relative expression of *Psen1*, *Bace1*, *App*, *Mapt*, and *Gsap* was normalized to *Gapdh* and expressed relative to CMC controls. **(C)** Western blot analysis of APP processing proteins. Representative immunoblots and densitometric quantification of APP, BACE1, PS1, and ADAM10, normalized to GAPDH, are shown. **(D)** Western blot analysis of phosphorylated ERK (pERK) normalized to GAPDH **(E)** Brain amyloid-β levels measured by ELISA; fold-change for Aβ40 and Aβ42 normalized to CMC are shown as indicated. Brains of SP-and SS-treated mice showed similar decreases in Adam10 and pERK, and a prominent increase in Aβ42 across all three treatment groups, suggesting that amyloidogenic alterations might not be attributable exclusively to the effect of SP on memory decline. Data are presented as mean ± SEM (n= 3-4 in each group); *p < 0.05, **p < 0.01, ***p < 0.001, ****p < 0.0001, One-way ANOVA, followed by Tukey’ multiple comparison test.

To determine whether these transcriptional changes were reflected at the protein level, we next assessed key APP-processing components by western blot. APP protein levels were reduced in SS-treated mice relative to CMC controls, whereas PS1 expression remained unchanged across groups (Fig. 4C). ADAM10 levels were significantly decreased in both SS-and SP-treated mice compared with controls (Fig. 4C). Consistent with this finding, phosphorylated ERK (pERK), a downstream signaling molecule linked to neuronal plasticity and regulation of the APP-processing pathway, was reduced across all bacterial treatment groups relative to CMC controls, with the greatest reduction observed in SS-treated mice (Fig. 4A and 4D).

Aβ production was next evaluated in brain lysates to assess the downstream consequences of the altered APP-processing pathway. Interestingly, ELISA analysis of brain lysates demonstrated Aβ40 levels were significantly increased in all NF-, SS-, and SP-treated mice relative to controls (Fig. 4E). Similarly, Aβ42 levels were likewise elevated in all bacterial treatment groups compared with CMC mice, with the most pronounced increase observed in NF-treated animals (Fig. 4E). Together, these unexpected findings indicate that chronic oral bacterial exposure shifts APP processing away from the non-amyloidogenic pathway, as evidenced by reduced pERK and ADAM10 expression, while promoting the accumulation of Aβ40 and Aβ42, consistent with a pro-amyloidogenic state. Aβ accumulation alone may not be sufficient to drive the cognitive deficits in this model.

### No evidence of oral *S. parasanguinis* translocation to the brain

Given the neuroinflammatory and neuropathological changes observed in SP-treated mice, we next investigated whether chronic oral exposure to SP was associated with bacterial translocation to the brain. Using an RNAScope-based SP-specific probe on sagittal FFPE mouse brain sections, no detectable SP-specific RNA signal was observed in any brain region (Fig. 5A). A positive control probe was included for fluorescent comparison and quantification (Fig. 5B).

**Figure 5.**
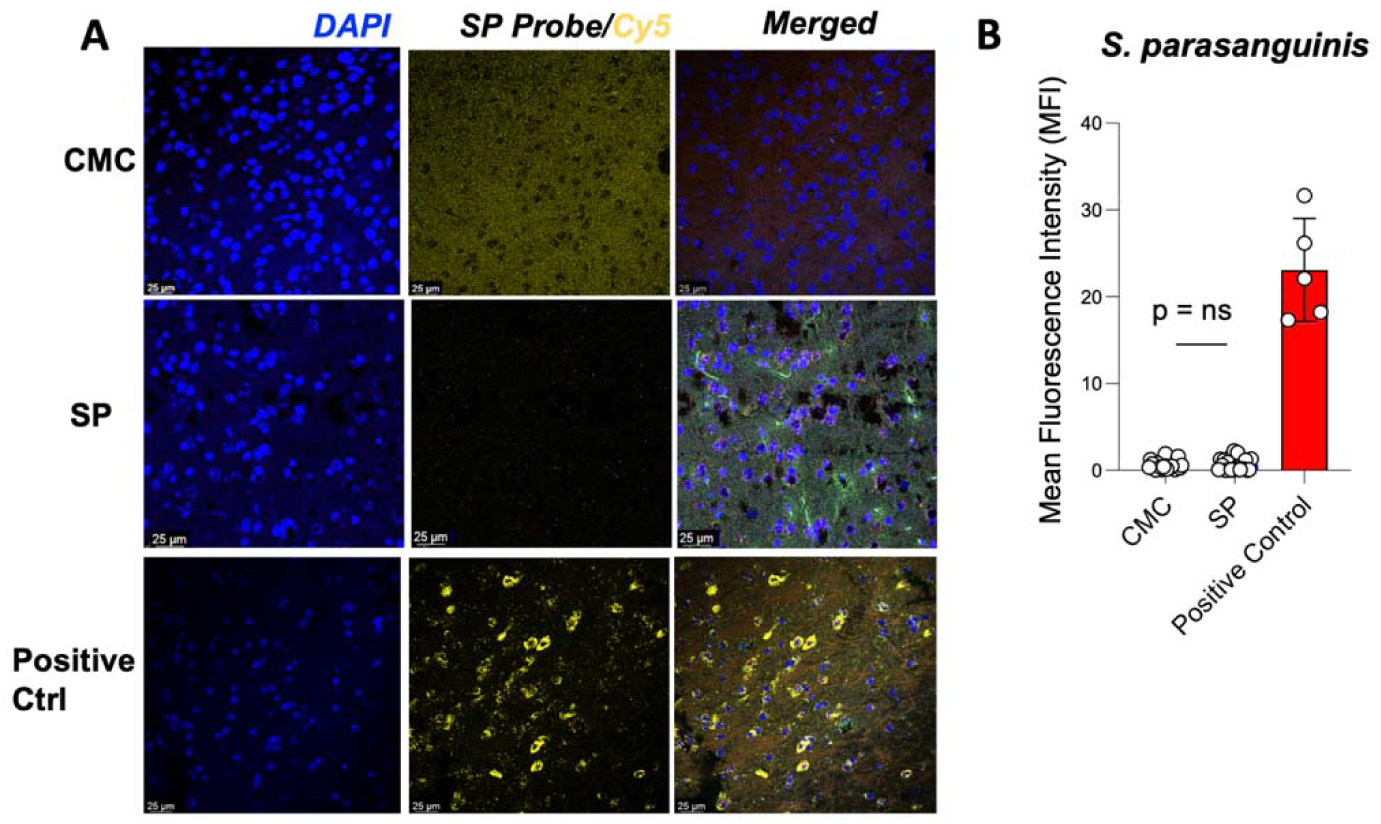
No evidence of *S. parasanguinis* translocation to the mouse brains. **(A)** Representative images obtained through RNAScope in situ hybridization of sagittal mouse brain sections (5 μm FFPE). Fluorescent imaging showed no detectable *S. parasanguinis* RNA signal (yellow, Cy5 channel) across the entire section. A positive probe (PPIB, a housekeeping gene) confirmed assay validity. Images were acquired at 40× magnification on a Zeiss LSM 510 Meta confocal microscope (1024 × 1024; n = 4). **(B)** Mean fluorescence intensity (MFI) wa calculated using ImageJ/Fiji, and bars indicate mean ± SEM.

### Untargeted metabolomics profiling of bacterial supernatants, mouse saliva, and brain identifies candidate bacterial metabolites

Recent advances in neuroscience have demonstrated that orally administered bacterial metabolites can rapidly penetrate the brain and peripheral tissues within 2 hours, with effects sustained for up to 18 hours ^41^. Given that the preceding experiments suggested that direct bacterial translocation to the brain was unlikely, we next investigated whether microbial metabolites might provide an indirect mechanistic link between oral bacterial exposure and altered brain function. Here, untargeted metabolomics identified 340 metabolites in mouse saliva, 919 in brain tissue, and 1682 in bacterial supernatants, revealing treatment-dependent alterations across multiple metabolite classes. Saliva metabolomic profiles were initially used to prioritize candidate metabolites. The top metabolites included compounds unique to SP-treated mice (cysteine S-sulfate [CSS] and maltotetraose), shared between SP-and SS-treated mice (histamine and sphingosine 1-phosphate), and metabolites common to all microbiome-treated groups (histidine betaine and N-acetylputrescine) (Fig. 6A). Cross-comparison with bacterial supernatant profiles demonstrated that SP and/or SS directly produced CSS, histamine, N-acetylputrescine, and related metabolites within these metabolic pathways, including sphingosine and cysteine (Fig. 6B).

**Figure 6.**
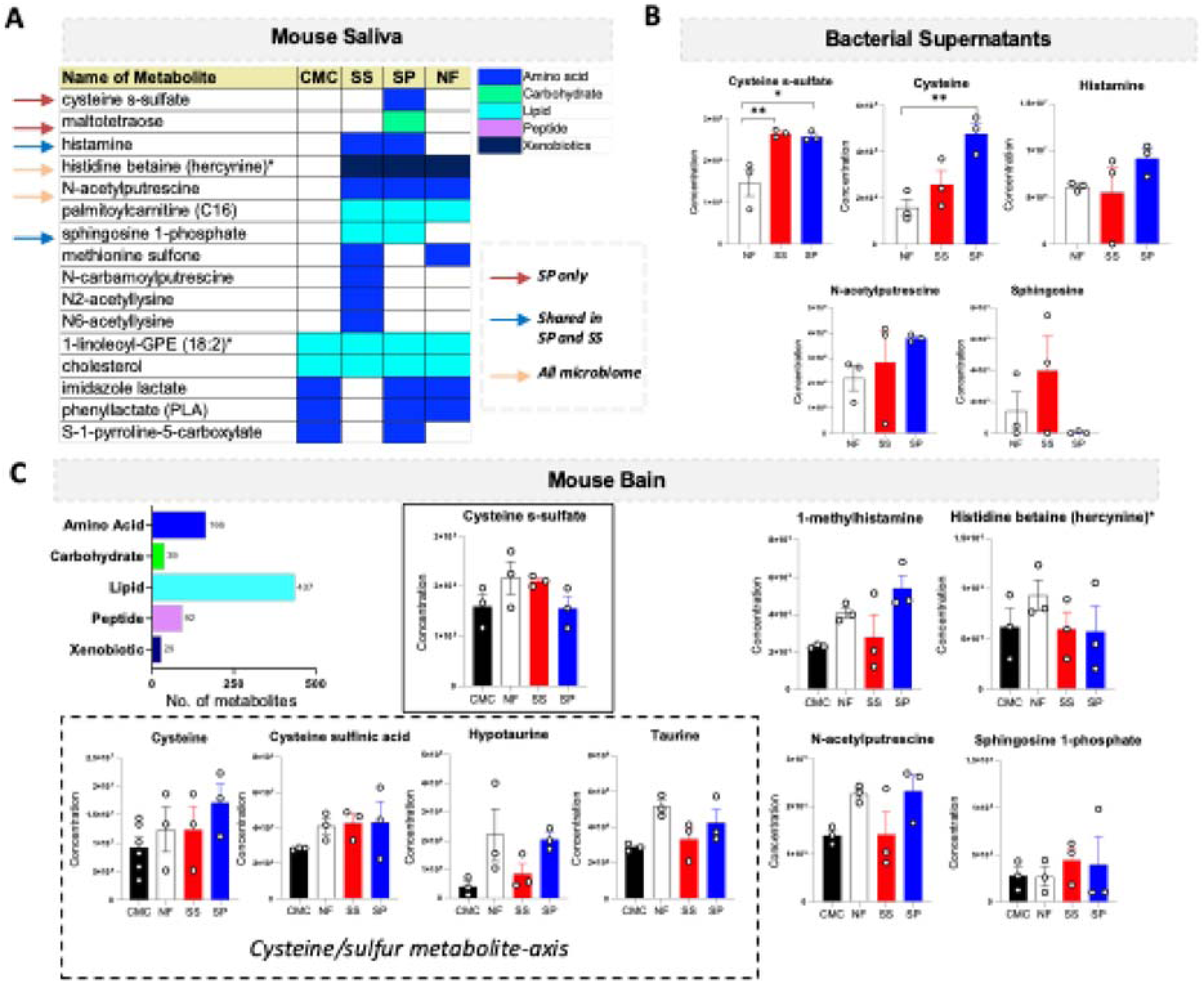
Untargeted metabolomic profiling of bacterial supernatants along with mouse saliva and brain following oral bacterial inoculations. **(A)** A detection matrix of salivary metabolites across treatment groups is shown. Colored cells indicate metabolites detected above threshold and grouped by biochemical class; blank cells denote non-detection. The top two metabolites (a total of six) were selected in each category, i.e., SP only, shared in SP and SS, and shared metabolite in all three microbiome treatment groups, for comparison with the other two metabolomic profiles. **(B)** Bacterial supernatants contained three exact metabolites (cysteine S-sulfate, histamine, and N-acetylputrescine) found in mouse saliva, as well as two chemically relevant metabolites, while maltotetraose and histidine betaine were not produced by the microbiome. **(C)** While screening the brain metabolomics, it appears that a chemically relevant metabolite of histamine (1-methylhistamine) was increased in the brains of SP-treated mice, though it was shared in the saliva of SP and SS, along with the modulation in the cysteine /sulfur metabolite axis in the SP-treated mouse brains (i.e., cysteine-s-sulfate, cysteine, cysteine sulfinic acid, hypotaurine, and taurine). Data are presented as mean ± SEM (n = 3 per group). *p < 0.05, **p < 0.01; one-way ANOVA with Tukey’s post hoc test.

We next examined whether these metabolites or metabolites related to metabolic pathways were detectable in the brain. Although CSS itself was not highly abundant in the brains of SP-treated mice, several metabolites associated with the cysteine/sulfur metabolic axis were elevated compared to the vehicle controls (Fig. 6C). In addition, 1-methylhistamine, a downstream metabolite of histamine, was significantly increased in the brains of SP-treated mice compared to controls (Fig. 6C). Other metabolites, including histidine betaine, N-acetylputrescine, and sphingosine 1-phosphate, were detected in both saliva and brain samples, although their brain abundance did not differ substantially between groups (Fig. 6C). Together, these findings suggest that SP uniquely produces or enriches CSS in the oral compartment, while inducing broader alternations in sulfur-amino-acid metabolism within the brain. This pattern supports the possibility that oral SP exposure perturbs cysteine/sulfur metabolic pathways rather than simply depositing a single metabolite unchanged into the brain. Furthermore, although histamine abundance was shared between both *Streptococcus* species in the oral compartment (Fig. 6A), the selective enrichment of histamine-related metabolites in SP-treated brains (Fig. 6C) suggests coordinated remodeling of cysteine/sulfur metabolism and histaminergic signaling during SP-associated neuroimmune responses.

### Cysteine-S-sulfate and histamine induce proinflammatory cytokine production in primary microglia and A**β** secretion *in vitro*

Given that CSS and histamine emerged as plausible SP-specific mechanistic candidates, we next evaluated their direct effects on neuroinflammatory and amyloid-associated pathways *in vitro*. In primary mouse microglia, CSS treatment induced a dose-dependent increase in IL-1β production, with higher concentrations also significantly increasing TNF-α levels (Fig. 7A). Histamine similarly elevated IL-1β and TNF-α production in microglia, supporting the possibility that SP-associated metabolites contribute to inflammatory signaling in the brain.

**Figure. 7:**
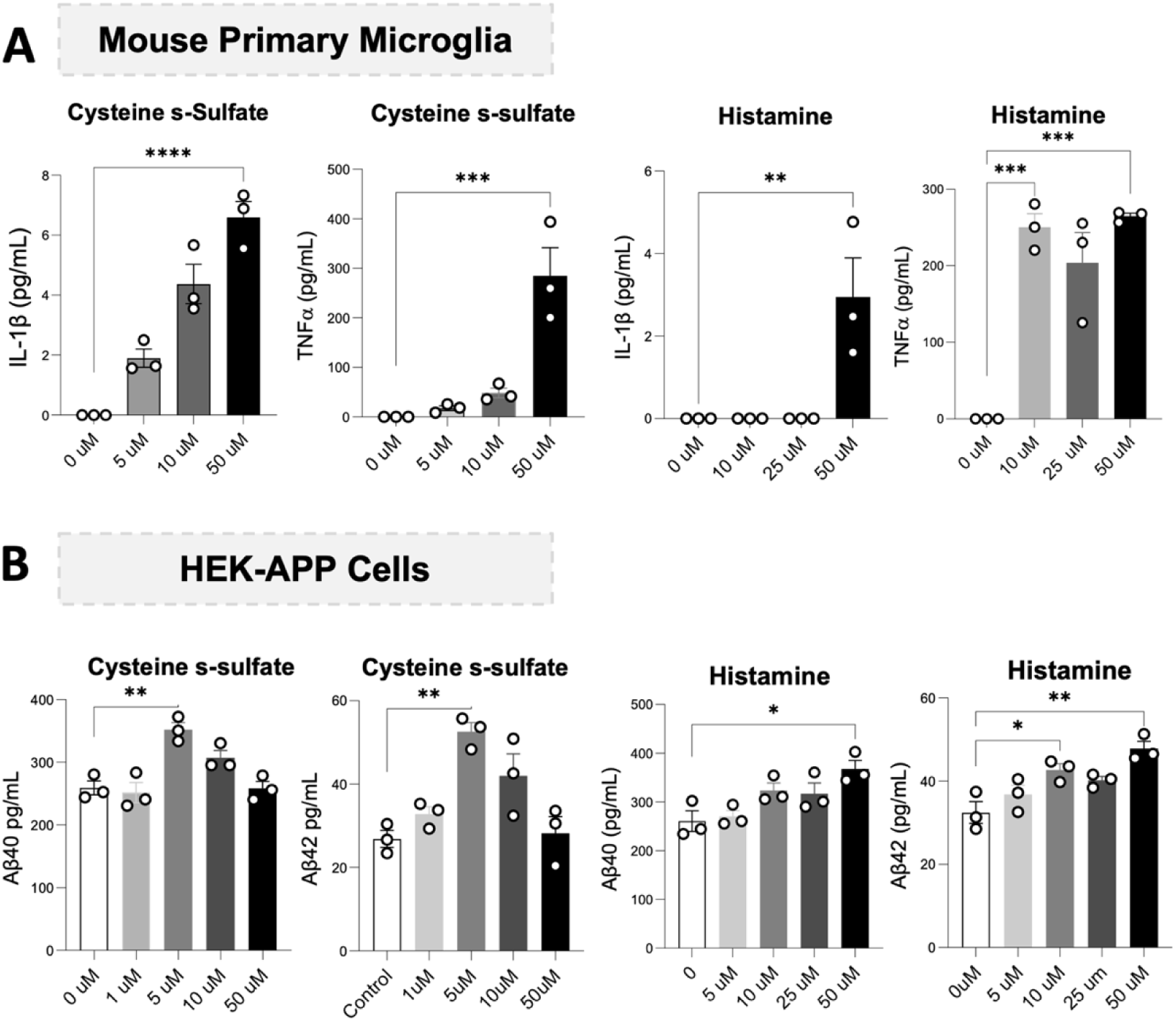
Metabolite-associated inflammatory and amyloidogenic responses in primary microglia and HEK-APP cells. **(A)** Increased IL-1β and TNF-α levels have been detected in supernatants from primary mouse microglia treated with indicated concentrations of cysteine s-sulfate and histamine. **(B)** Aβ40 and Aβ42 levels were increased in response to cysteine s-sulfate and histamine at the indicated concentrations in HEK-APP cells in vitro. The assays were run in triplicate. Data are presented as mean ± SEM (n = 3 per group); *p < 0.05, **p < 0.01, ***p < 0.001, ****p < 0.0001, One-way ANOVA, followed by Tukey’s multiple comparison test.

We next examined the direct effect of these metabolites on amyloid-associated pathways using HEK-APP cells *in vitro*. CSS treatment significantly increased Aβ40 and Aβ42 production at intermediate concentrations compared to untreated controls (Fig. 7B). Histamine treatment also modestly increased both Aβ40 and Aβ42 production in a dose-dependent effect. In contrast, several additional candidate metabolites, including N-acetyl-putrescine and sphingosine-1-phosphate, enriched in the oral cavity of all bacterial groups (Fig. 6A), and taurine (a neuroprotective metabolite control, ^42^), did not alter Aβ40 and Aβ42 production at any tested concentration compared to the untreated control (Suppl. Fig. 5). Moreover, to determine whether CSS alters microglial amyloid clearance capacity, human iPSC-derived microglia were treated with CSS for 24 hours, followed by flow cytometry analysis of Aβ42 phagocytosis. CSS treatment did not significantly affect microglial uptake of Aβ42 compared to untreated controls, suggesting that CSS-associated inflammatory responses occur independently of major alterations in Aβ42 phagocytic activity in microglia (Suppl. Fig. 6). Together, these findings suggest that SP-associated metabolites may contribute to neuroinflammatory and amyloid-related signaling pathways relevant to SP-associated neuropathogenesis.

## Discussion

Antibiotic treatment alters behavioral responses to cocaine, indicating a role for microbiota in cocaine addiction ^43^. Emerging evidence links oral microbial dysbiosis, particularly chronic periodontitis, to increased brain Aβ and risk of AD in humans ^44,45^. The periodontal pathogen *Porphyromonas gingivalis* (*Pg*) and its gingipains are detected in the brains of AD patients; *Pg* oral infection increases tau levels in the brains of wild-type C57BL/6 mice ^46^. Aβ aggregation, an early AD hallmark ^47^, is observed in mice with oral *Pg* infection, alongside brain bacterial colonization ^46^. However, cognitive decline following *Pg* treatment manifests only when additional risk factors, such as aging or high-fat diet-induced obesity, are present ^46,48–50^. In the present study, SP induced memory impairment in wild-type B6 mice aged 20 weeks, earlier than the cognitive decline observed in transgenic AD models (∼27 weeks ^51,52^), highlighting its neurodegenerative potential.

Prior studies have demonstrated that chronic cocaine use is associated with oral microbial dysbiosis and immune perturbation ^28^. Our previous human study further identified enrichment of SP in saliva samples from CUD ^19^, and the current study demonstrated that cocaine directly promoted SP growth *in vitro*. Building on these observations, we investigated whether oral administration of cocaine-enriched bacteria in wild-type B6 mice could elicit behavioral and neuropathological changes relevant to cocaine-associated neurobiological disorders. In the present study, oral inoculation of SP for three months resulted in spatial memory deficits, microglial activation, and neuroinflammation in wild-type B6 mice. Although no evidence of bacterial translocation to the brain was detected, SP-produced metabolites, including CSS and histamine, were enriched in the saliva of SP-treated mice and were capable of inducing inflammatory responses in microglia and promoting Aβ secretion *in vitro*.

Mice treated with SP exhibited significant impairments in spatial learning and memory, as measured by the Barnes maze. Importantly, these effects occurred in the absence of detectable bacterial infection or translocation in the brain, reinforcing the hypothesis that indirect mechanisms, such as metabolite-mediated signaling, may underlie the observed neuroinflammatory and neurobiological alterations. Metabolomic profiling further supported this interpretation, as SP-treated mice exhibited distinct metabolite signatures characterized by enrichment of CSS and altered histamine-related metabolites, suggesting potential involvement of sulfur-associated metabolic pathways and histaminergic signaling in SP-associated neuropathogenesis. Notably, inflammatory responses were more prominent in the brain than in peripheral tissues, suggesting dose-dependent oral-to-brain penetration of metabolites and preferential engagement of neuroimmune pathways compared to the periphery. Consistent with this interpretation, non-region-specific microglial activation may contribute to increased IL-1β production and Aβ accumulation, both of which are well-established features of early neurodegenerative processes ^53–57^.

Aβ42 accumulation is widely recognized as a key feature of AD and is closely linked to amyloid plaque formation in the brain ^58,59^. Unexpectedly, all bacterial treatment groups exhibited elevated Aβ42 levels, with the greatest increase observed in NF-treated mice. Moreover, SP treatment was accompanied by transcriptional and protein-level changes consistent with altered APP processing, including reduced *Bace1/Psen1* transcript expression, increased *Gsap* expression, reduced ADAM10, and decreased pERK signaling. These findings are particularly notable because ADAM10 is a critical mediator of the non-amyloidogenic APP-processing pathway, whereas reductions in ERK-linked signaling may reflect impaired neuroprotective or synaptic regulatory mechanisms ^60,61^. Together, these findings suggest that chronic exposure to cocaine-associated oral bacteria may modulate APP-processing and amyloid-associated signaling pathways in the brain. Reduced ADAM10 and pERK expression suggests suppression of the non-amyloidogenic α-secretase pathway. Although canonical amyloidogenic markers (BACE1 and PSEN1) were not significantly elevated, increased Aβ40 and Aβ42 levels indicate a shift in APP metabolism toward a more amyloidogenic state.

While amyloidogenic pathways appear to be dysregulated across bacterial treatment groups, only SP-treated animals exhibited significant impairments in spatial learning and memory, suggesting that amyloid accumulation alone may be insufficient to drive cognitive decline. Instead, elevated IL-1β observed in the brain of SP-treated mice may play a more central role by acting in concert with amyloid-related changes to impair synaptic function. IL-1β is a potent pro-inflammatory cytokine with well-established effects on synaptic plasticity, particularly in the hippocampus, a region critical for spatial learning and memory ^62–64^. Chronic elevation of IL-1β impairs long-term potentiation and disrupts excitatory NMDA signaling ^65–67^. Beyond its direct effects on synaptic function, increased IL-1β may also contribute to microglial activation and maladaptive synaptic remodeling. Because excitatory synapses are concentrated on dendritic spines, sustained neuroinflammatory signaling may reduce synaptic connectivity and disrupt neural integration in regions important for spatial learning and memory ^62,64,65^. These mechanisms provide a plausible link between elevated brain IL-1β levels and the spatial memory deficits observed in SP-treated mice. Importantly, inflammatory and metabolite-associated mechanisms are not mutually exclusive and may converge to disrupt synaptic integrity and neuronal signaling.

Microbial-derived metabolites, such as CSS, may further exacerbate synaptic dysregulation through excitotoxic glutamatergic signaling pathways ^21,68–70^. In contrast to IL-1β-mediated inflammatory disruption of synaptic plasticity, CSS may promote neuronal injury by overstimulating glutamatergic signaling, thereby increasing calcium influx, oxidative stress, and mitochondrial dysfunction, all of which are characteristic features of excitotoxic damage ^71–74^. Additionally, the increased abundance of 1-methylhistamine in SP-treated brains may reflect altered histamine turnover and dysregulated histaminergic signaling. Beyond its role as a neurotransmitter, histamine can modulate microglial activation and cytokine production, and excessive or dysregulated histaminergic activity has been associated with neuroinflammation and impaired cognitive function ^75^.

In the present study, CSS was uniquely enriched in saliva from SP-treated mice, detected in bacterial supernatants, and demonstrated bioactivity *in vitro* by inducing microglial cytokine release and altering Aβ-associated pathways. Although CSS itself was not prominently elevated in the brain metabolomic dataset, SP-treated brains exhibited broader alterations in chemically related sulfur-associated metabolites, including metabolites linked to cysteine and taurine metabolism, suggesting engagement of a cysteine/sulfur metabolic axis rather than simple accumulation of a single metabolite within the brain. It is also important to note that untargeted metabolomics approaches may fail to detect low-abundance, transient, or spatially restricted metabolites in brain tissue ^76,77^. In parallel, increased levels of the histamine-related metabolite 1-methylhistamine in SP-treated brains suggest further remodeling of histaminergic signaling pathways, which have been implicated in neuroinflammatory and cognitive processes ^75^. Previous studies have demonstrated that microbial metabolites can reach the brain within 2 hours of oral administration ^41^, supporting a potential role for SP-associated metabolites in oral-brain communication. However, indirect mechanisms mediated by peripheral immune modulation or systemic inflammatory signaling cannot be excluded ^78,79^. Taken together, these findings support CSS and altered histamine-associated metabolites as plausible contributors to SP-associated neuroimmune effects while underscoring the need for future studies to define the mechanisms underlying microbiome-mediated oral-brain signaling.

A limitation of the present study is that the causal contribution of individual downstream mediators (e.g., CSS) *in vivo* remains unresolved. Although CSS and altered histamine-associated metabolites emerged as candidate mediators of SP-associated effects, direct testing in animal models will be required to determine whether they are sufficient to reproduce the observed neuroimmune and cognitive effects *in vivo*. In addition, although elevated brain IL-1β was associated with SP-induced memory impairment, future studies are needed to determine whether IL-1β signaling directly causes the observed neurobehavioral phenotype without cooperating with Aβ42 or other neurodegeneration pathologies. Because bacterial localization was not detected in the brain and the current analyses relied primarily on whole-brain metabolomics and selected imaging approaches, the precise route and regional specificity of oral SP-associated CNS effects also remain unclear. Future investigations should therefore determine whether CSS and histamine-associated signaling preferentially affect memory-related brain regions and whether these effects can be modified through microbial depletion, metabolite-signaling blockade, or anti-inflammatory interventions. Finally, the mechanisms through which cocaine selectively enhances SP growth *in vitro* remain unknown and warrant further investigation, including whether SP directly metabolizes cocaine or instead adapts to cocaine-induced alterations in the oral microenvironment.

## Conclusion

This study demonstrates that chronic oral exposure to *S. parasanguinis* (SP) can promote neuroimmune activation and selective cognitive impairment. SP-treated mice exhibited microglial activation, elevated brain IL-1β, altered APP/amyloid-associated signaling, and deficits in spatial learning, while metabolomic profiling identified distinct microbial metabolites, including cysteine S-sulfate (CSS) and altered histamine-associated metabolites, as candidate mediators of SP-associated neuroimmune effects. Together, these findings support a model in which perturbations of the oral microbiota interact with neuroimmune and metabolic pathways to influence cognitive function. By identifying SP-associated inflammatory and metabolic alterations linked to memory impairment, this work highlights the oral-brain axis as a previously underrecognized contributor to chronic cocaine use-associated neurodegeneration.

## Materials and Methods

### Bacterial growth curve assay

*S. parasanguinis* strain F0449 (HM-808; BEI Resources, Manassas, VA) was cultured overnight in Trypticase Soy Broth (TSB; Sigma-Aldrich, St. Louis, MO) at 37 °C, diluted to OD600 = 0.1 in ZMB1 chemically defined medium ^19^, and supplemented with 100 mM glucose. Bacterial cultures were treated with cocaine hydrochloride (5 and 50 µg/mL; Sigma-Aldrich) or medium control. The lower dose of cocaine reflects upper-range physiological blood levels ^19^. Growth in triplicate was monitored by OD600 over 20 h at 37°C ^19^. Additional strains included *S. thermophilus* (BAA-491; ATCC, Manassas, VA) and *S. australis* (DSM 15627; DSMZ, Braunschweig, Germany).

### Experimental animals

C57BL/6J mice were obtained at 4-6 weeks of age from Jackson Laboratories (Bar Harbor, ME) and housed in the ABSL-2 facility under pathogen-free conditions with access to food and water *ad libitum*. All animal procedures were approved by the Medical University of South Carolina (MUSC Institutional Animal Care and Use Committee, IACUC protocol #IACUC-2019-00858-1), and all experiments were conducted in accordance with the Guide for the Care and Use of Laboratory Animals. Mice were allowed to acclimate for 7 days before the study began.

### Mouse study design and bacterial administration

Male and female mice were randomly assigned to one of four treatment groups (n = 5 per sex per group): *Streptococcus parasanguinis* (SP), *Streptococcus salivarius* (SS), *Neisseria flavescens* (NF), or vehicle control (2% carboxymethylcellulose [CMC]). Prior to bacterial administration, all mice, including CMC controls, received a 5-day course of oral antibiotics in the drinking water containing amoxicillin and vancomycin (1 mg/mL each) to reduce the native oral microbiota. Bacterial strains were grown to mid-log phase and prepared at a final concentration of 1 × 10□ cells/mL in sterile 2% CMC. A total volume of 90 µL of inoculum was administered directly into the oral cavity, with animals restrained according to protocol, twice weekly for 12 weeks. SP (strain F0449, HM-808) and NF (strain SK114, HM-115) were obtained through BEI Resources (Manassas, VA), NIAID, NIH, as part of the Human Microbiome Project. SS (strain BAA-1024™) was obtained from the American Type Culture Collection (ATCC, Manassas, VA, USA). Body weight was monitored weekly throughout the study. At the study endpoint, oral samples were collected for microbiological confirmation of colonization, and tissues and biofluids were harvested for downstream molecular, biochemical, and metabolomic analyses.

### Biochemical confirmation of colonization

To confirm successful colonization and distinguish between bacterial strains, oral samples were cultured on chocolate blood agar (for NF – Hardy Diagnostics, Santa Maria, CA, Cat. No. E14) and *Streptococcus*-selective media (for SP and SS – Hardy Diagnostics, Santa Maria, CA, Cat. No. A70). Colonies were evaluated via Gram staining, catalase testing, and hemolytic activity on blood agar. Urease activity was assessed using Christensen’s urea agar slants ^80^. Urease-positive SS colonies turned the slant pink, whereas SP remained urease-negative and produced no color change. Representative colonies were visualized by light microscopy.

### Behavioral tests

All behavioral experiments were conducted in the MUSC Behavioral Core during the light phase (between 9 AM and 5 PM). Mice were allowed to habituate to the testing room for at least 30 min before each session. Behavioral activity was recorded and analyzed using ANY-maze software (Stoelting Co, Chicago, IL, USA). All testing equipment was cleaned with 70% ethanol between sessions to minimize odor cues.

### Open field test

The open field test (OFT) was performed to assess general locomotor and exploratory behavior. Mice were placed in a square arena and allowed to freely explore for 10 min. Total distance traveled was used as the primary outcome measure.

### Elevated plus maze

The elevated plus maze (EPM) was used to assess anxiety-like behavior. The apparatus consisted of two open arms, two enclosed arms, and a central starting area. Each mouse was placed in the center of the maze and allowed to explore for 5 min. Time spent in the open and closed arms, as well as the number of arm entries, was recorded.

### Novel object recognition

The novel object recognition (NOR) task was used to assess recognition memory. Mice were first habituated to the testing arena, followed by a familiarization phase in which they were exposed to two identical objects. During the testing phase, one familiar object was replaced with a novel object. Time spent interacting with the familiar and novel objects was recorded, and the discrimination index was calculated as (time spent with the novel object minus time spent with the familiar object) divided by total exploration time.

### Barnes maze

The Barnes maze was used to assess spatial learning and memory. The apparatus consisted of an elevated circular platform containing 20 holes, one of which led to an escape chamber. Distal visual cues were positioned around the maze to facilitate spatial navigation. Mice underwent training trials over five consecutive days, followed by a test trial to evaluate memory retention. Outcome measures included test duration, latency to first reach the escape hole, latency to first enter the escape chamber, time spent in the escape hole zone, time spent in the escape quadrant, and entries into the escape quadrant. Heat maps and representative track plots were generated via ANY-Maze software to qualitatively assess the search strategy.

### Forced swim test

The forced swim test (FST) was performed to assess stress-coping-related behavior. Mice were placed in a transparent cylinder filled with water for 6 min, and the total immobility and mobility times were recorded.

### Brain and tissue collection

At study completion, mice were euthanized, and brains were rapidly extracted and hemisected. One hemisphere was snap-frozen for downstream RNA, protein, ELISA-based, and metabolomic analyses, while the other hemisphere was immersion-fixed in 4% paraformaldehyde for RNAscope in situ hybridization. Fixed tissue was paraffin-embedded according to standard procedures, sectioned, and mounted on charged slides for imaging.

### RNAscope in situ hybridization

RNAscope in situ hybridization was performed using the RNAscope Multiplex Fluorescent V2 assay (ACD Bio-Techne, Newark, CA) according to the manufacturer’s instructions. Formalin-fixed paraffin-embedded brain sections were pretreated, including target retrieval, followed by probe hybridization and signal amplification. Probes were used to detect Iba1 (catalog no. 1089911-C1), Tmem119 (catalog no. 472901-C4), universal bacterial 16S rRNA (catalog no. 464461-C4), a SP-specific sequence (catalog no. 1633621-C2), and the PPIB standard positive control probe (housekeeping gene). DAPI was used as a nuclear counterstain. Fluorescent images were acquired using a Leica Stellaris 8 confocal microscope. Quantification of fluorescence signal was performed in Fiji/ImageJ using standardized analysis parameters applied across all groups.

### RNA isolation, cDNA synthesis, and quantitative PCR

RNA was extracted from frozen whole-brain lysates using TRIzol reagent per the manufacturer’s protocol. cDNA synthesis was performed using the SuperScript III First-Strand Synthesis System (Invitrogen, Carlsbad, CA). Gene expression was quantified using TaqMan™ Fast Advanced Master Mix (Cat No. 4444557; Thermo Fisher Scientific, Waltham, MA) and pre-designed TaqMan probes on a Bio-Rad CFX Connect system. Reactions were run in 96-well plates using 15 µL volumes, with standard cycling conditions (95°C for 10 min, followed by 40 cycles of 95°C for 15 s and 60°C for 30 s).

Quantitative PCR (qPCR) was performed using TaqMan® Gene Expression Probes (Thermo Fisher Scientific, Waltham, MA) targeting key inflammatory and neurodegenerative genes. Probes included IL-6 (Mm00446190_m1), TNF-α (Mm00443258_m1), and IL-1β (Mm00434228_m1) for inflammation profiling. Neurodegenerative markers assessed were APP (Mm01344172_m1), MAPT (Mm00521988_m1), BACE1 (Mm0478664_m1), GSAP (Mm00615236_m1), and PSEN1 (Mm00501184_m1). Gapdh (Mm99999915_g1) served as the endogenous control. Reactions were run according to manufacturer protocols, and relative gene expression was calculated using the ΔΔCt method.

### Meso scale discovery (MSD) multiplex cytokine assay

Pro-inflammatory cytokines were quantified in lysed brain tissue, serum, gut, and saliva using the U-PLEX Custom Biomarker Group Assay (MSD, Rockville, MD). Sample preparation and plate loading were carried out following the manufacturer’s instructions. Plates were read on a MESO QuickPlex SQ 120 reader, and concentrations were calculated using Discovery Workbench software. The number of biological replicates varied across assays (n = 3-5 per group), with all available samples included in each analysis.

### Enzyme-linked immunosorbent assays (ELISAs)

Corticosterone levels in serum were measured using the Abcam Corticosterone ELISA Kit (Abcam, ab323743, Waltham, MA) following the manufacturer’s protocol. Aβ42 and Aβ40 levels were quantified in brain lysates using Invitrogen ELISA kits (Aβ42: Cat No. KMB3441; Aβ40: Cat No. KMB3481), with samples prepared in RIPA buffer containing protease inhibitors and normalized by total protein content determined via BCA assay. For in vitro experiments, HEK293 cells stably expressing human APP (AcroBiosystems, Newark, DE) were treated with different concentrations of cysteine-S-sulfate (CSS, Cat No. C2196-5MG; Sigma-Aldrich, Burlington, MA), and Aβ42 secretion was quantified using the same Invitrogen KMB3441 ELISA kit.

### Western blot analysis

Whole mouse brain tissues were washed with PBS and homogenized in RIPA lysis buffer containing 25 mM Tris-HCl (pH 7.6), 150 mM NaCl, 1% Nonidet P-40, 1% sodium deoxycholate, and 0.1% SDS, supplemented with protease and phosphatase inhibitors (Sigma-Aldrich, Cat No. PPC2020). Homogenates were incubated on ice for 30 minutes, then centrifuged at 12,000 × g for 15 minutes at 4 °C, and the supernatants were collected. Protein concentrations were determined using a BCA assay. Equal amounts of protein (20 μg per well) were separated by SDS-PAGE on 4-12% gels and transferred to PVDF membranes. Membranes were blocked in 5% milk in TBS-T for 1 hour and incubated overnight at 4 °C with primary antibodies against APP, BACE1, PS1-CTF, ADAM10, phospho-ERK, and GAPDH. The APP antibody was purchased from Abcam, BACE1, PS1-CTF, phosphor-ERK, and GAPDH antibodies were purchased from Cell Signaling Technology (Danvers, MA), and the ADAM10 antibody was purchased from MilliporeSigma (Burlington, MA). After washing, membranes were incubated with appropriate HRP-conjugated secondary antibodies and visualized using enhanced chemiluminescence substrate. Blots were imaged using a ChemiDoc imaging system, and band intensities were quantified using ImageJ. Target protein expression was normalized to GAPDH and expressed as a fold change relative to control samples.

### Metabolomic profiling

Saliva, whole-brain, and bacterial culture supernatant samples were submitted to Metabolon, Inc. (Durham, NC, USA) for untargeted metabolomic profiling. Samples were stored at −80 °C until analysis. Metabolites were extracted using methanol-based protein precipitation and analyzed by ultra-high-performance liquid chromatography tandem mass spectrometry (UPLC-MS/MS) using multiple chromatographic methods with positive and negative electrospray ionization to maximize metabolite coverage. Metabolites were identified by comparison to authenticated standards in Metabolon’s reference library based on retention index, accurate mass match, and MS/MS spectral similarity. Peak areas were quantified as the area under the curve. For visualization of saliva and brain metabolite detection matrices, metabolites were classified as detected or not detected across treatment groups based on the analytical output returned by Metabolon. For bacterial culture supernatants, clarified supernatants were collected after culture, filtered, and analyzed using the same untargeted metabolomic platform. Absolute peak intensity values for selected metabolites, including CSS, were used for downstream comparisons.

### In vitro cell treatment and neuroinflammation/neuropathogenesis quantification Chemicals

L-cystine S-sulfate, histamine, and sphingosine were purchased from Sigma. N-acetylputrescine hydrochloride was purchased from ChemCruz (Dallas, TX).

### Cell culture

HEK293T cells stably overexpressing human APP695 were cultured in Dulbecco’s modified Eagle’s medium at 37°C and 5% CO_2_. Cell lines were grown in this medium supplemented with 10% fetal calf serum. CSS, sphingosine, N-acetyl putrescine, and histamine were added for 24 h in HEK-APP before harvest.

### Primary mouse microglia

Primary mouse microglia derived from C57BL/6 mice (Creative Biolabs, Shirley, NY) were cultured according to the manufacturer’s instructions. For cytokine measurements, microglia were plated at a density of 0.5–1 × 10□ cells/cm² and allowed to adhere overnight. Cells were then treated with CSS for 24 h. Control cells received vehicle treatment. Levels of IL-1β and TNF-α in the culture medium were quantified using commercial mouse-specific ELISA kits (Abcam), according to the manufacturer’s instructions.

### Differentiation of human iPSC-derived microglia

Human iPSC-derived microglia were generated using a commercial stem cell differentiation kit according to the manufacturer’s protocol. Differentiated microglia were maintained in microglia maturation medium and used for experiments after achieving a mature microglial phenotype.

### Flow cytometry analysis of microglial phagocytosis for A**β**42

Phagocytic activity of human primary microglial cells was assessed using a TAMRA-labeled amyloid-β (Aβ□□□□) peptide (AnaSpec, Fremont, CA). Primary microglia were plated at the indicated density and treated with CSS for 24 h. Following treatment, cells were incubated with TAMRA-conjugated Aβ□□□□(1 μM) for 1 h at 37 °C to allow peptide uptake. After incubation, cells were washed extensively with cold PBS to remove unbound extracellular peptide and detached using Accutase (Stemcells, Newark, CA). Cells were collected, resuspended in flow cytometry buffer (PBS containing 2% fetal bovine serum), and analyzed using a flow cytometer. TAMRA fluorescence was detected in the appropriate fluorescence channel, and phagocytic activity was quantified as the percentage of TAMRA-positive cells and/or mean fluorescence intensity (MFI). Data were analyzed using FlowJo software. Background fluorescence was determined using cells not exposed to TAMRA-Aβ and was subtracted from experimental samples.

### Quantification of A**β**42 and A**β**40

Levels of Aβ42 and Aβ40 in both the culture medium and mouse brain lysate were measured in triplicate using sandwich ELISA kits from Thermo Fisher Scientific according to the manufacturer’s instructions.

### Quantification of IL-1**β** and TNF-**α**

IL-1β and TNF-α levels were measured in triplicate in cell culture supernatants using mouse ELISA kits (Abcam) following the manufacturer’s protocol.

## Statistical analysis

Statistical analysis was performed using GraphPad Prism 10 software (GraphPad Software, San Diego, CA). Data are represented as mean ± SEM, as indicated in the corresponding figure legends. For comparisons among treatment groups (CMC, SP, SS, and NF), one-way and two-way analyses of variance (ANOVAs) were used, followed by Tukey’s multiple comparisons test when appropriate. A p-value of < 0.05 was considered statistically significant.

## Acknowledgment

`o` Behavioral testing was supported by the CNDD Mouse Behavior Phenotyping Core (P20GM148302), a University Shared Research Facility at MUSC.
`o` Image facilities were supported in part by the Cell & Molecular Imaging Shared Resource, MUSC Cancer Center Support Grant (P30 CA138313), the SC COBRE in Digestive and Liver Diseases (P20 GM130457), the MUSC Digestive Disease Research Cores Center (P30 DK123704), and the Shared Instrumentation Grants S10 OD018113 and S10 OD028663.
`o` Immunohistology work was supported in part by the Translational Science Shared Resource, Hollings Cancer Center, Medical University of South Carolina (P30 CA138313).

## Author Contributions

Conceptualization: WJ, DJ

Methodology: DJ, TS, BB, HS, AN, KS

Investigation: DJ, TS, AN, BB, ZL

Data analysis and Visualization: DJ, TS, AN, ZW

Supervision: WJ, TS

Funding acquisition: WJ

Writing-original draft: DJ, TS

Writing-reviewing & editing: DJ, TS, WJ, RDP, AN

## Data and material availability

All data are available in the main text or the supplementary materials.

The raw files regarding the behavioral assessments and metabolomic screening in mice saliva and brain have been deposited and can be found at: https://doi.org/10.6084/m9.figshare.32399916

## SUPPLEMENTAL DATA/FIGURES

**Supplemental Figure 1.**
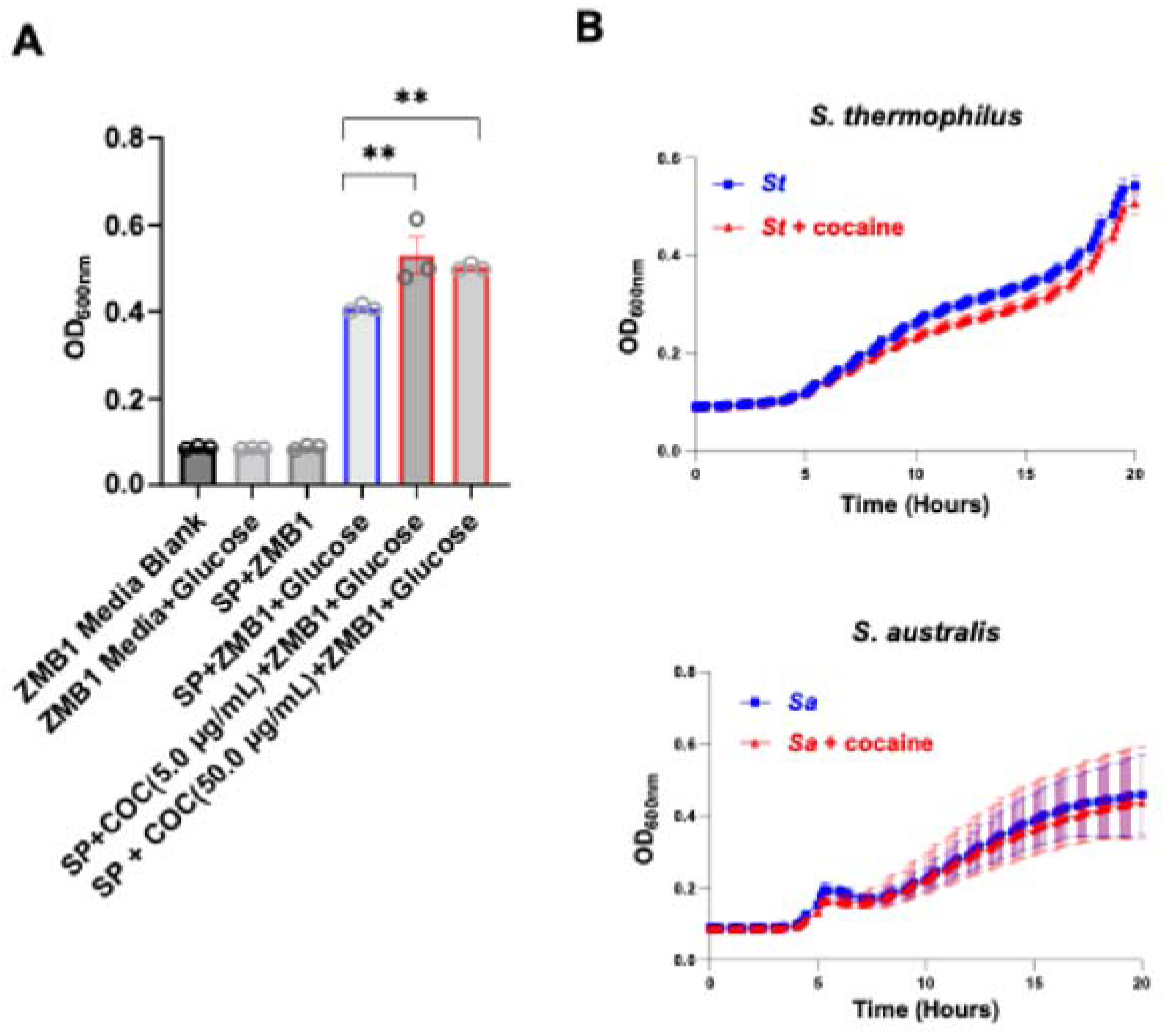
Cocaine does not promote the growth of *S. australis* or *S. thermophilus* in vitro. **(A)** Dose-response curve for the *S. parasanguinis* in the presence of cocaine (5 and 50 ug/mL) in ZMB1 media with glucose (50mM). **(B)** *S. australis* and *S. thermophilus* were also enriched in CUD (Johnson et al., BioRxiv 2025); however, they showed no significant increase in growth curve pattern in the presence of cocaine (5 ug/mL), suggesting cocaine targets *S. parasanguinis* specifically. Bacterial growth curve was assessed by Optical density (OD600). Data are presented as mean ± SEM (triplicate); **p < 0.01, One-way ANOVA, followed by Tukey’s multiple comparison test.

**Supplemental Figure 2.**
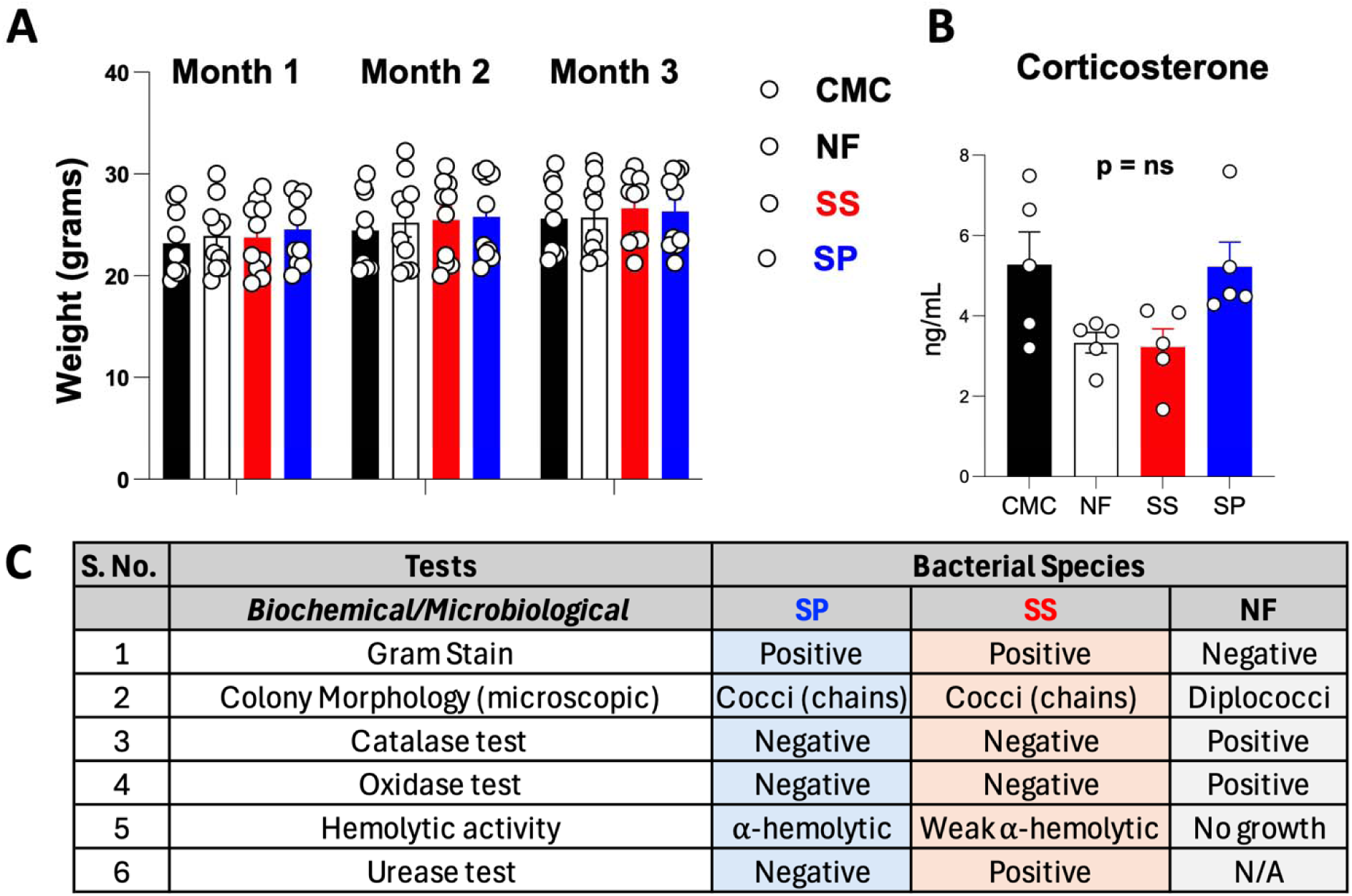
No change in body weight and serum corticosterone levels in mice. **(A)** Baseline assessments before behavioral testing. Body weight increased normally across groups during chronic oral inoculation with *S. parasanguinis, S. salivarius,* and *N. flavescens*. **(B)** Serum corticosterone levels remained unchanged, indicating no handling-or environment-related stress. Data are presented as mean ± SEM (n = 5-10 per group), One-way ANOVA, followed by Tukey’s multiple comparison test. **(C)** Microbiological and biochemical characterization of bacteria recovered from the oral cavity at the study endpoint. Recovered isolates were evaluated by Gram staining, colony morphology, catalase testing, oxidase testing, hemolytic activity, and urease testing, confirming species-consistent profiles for *S. parasanguinis, S. salivarius,* and *N. flavescens* following repeated oral administration.

**Supplemental Figure 3.**
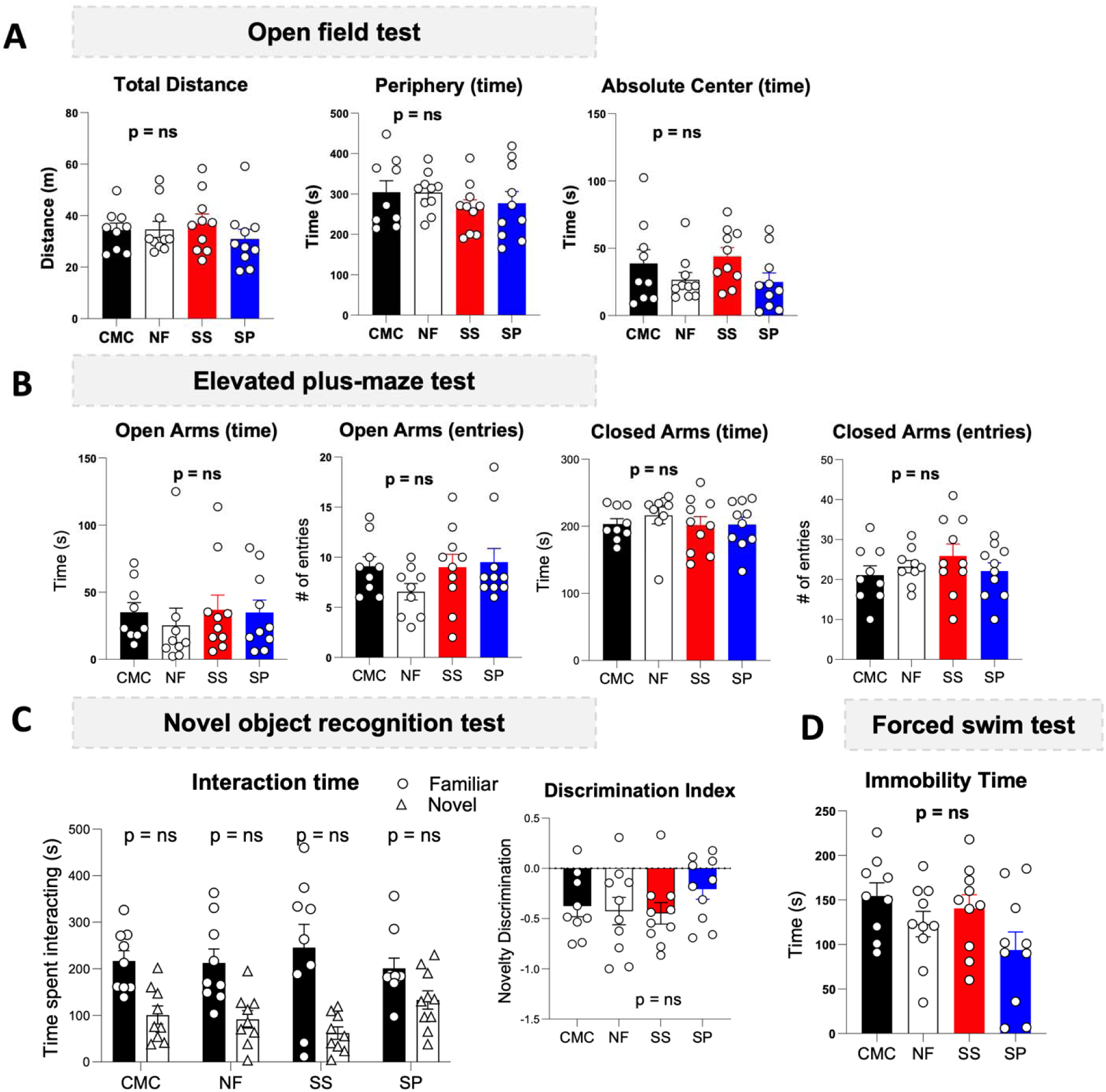
No significant changes in mouse behaviors related to recognition memory, anxiety, and depression. **(A)** Quantification of open field test (OFT) parameters, i.e., no significant change in total distance covered (F*_(3,35)_*= 0.2502, p =0.86), time spent in the periphery (F*_(3,35)_* = 1.872, p =0.15), and the time spent in the absolute center (F*_(3,35)_* =1.056, p= =0.38) in SP-treated mice as compared to controls. **(B)** Different parameters monitored in the Elevated plus-maze (EPM) test, showing no significant differences in time spent in open arms(F*_(3,34)_* = 0.104, p = 0.95), and closed arms F*_(3,34)_* = 0.653, p = 0.58) as well as number of entries in open arms F*_(3,34)_* = 0.287, p = 0.83) and closed arms F*_(3,34)_* = 1.13, p = 0.34) in by SP-treated mice compared to controls. **(C)** The Novel object recognition (NOR) test showed no significant impairment in recognition memory, as indicated by the total interaction time with familiar versus novel objects (F*_(3,66)_* = 1.65, p =0.18) and the discrimination index (F*_(3,35)_* = 0.695, p = 0.56) in SP-treated mice compared to controls. **(D)** Forced swim test (FST) also showed no significant differences among the groups in immobility time (F*_(3,35)_* = 0.475, p = 0.70). Data are presented as mean ± SEM (n = 9-10 per group). One-way and two-way ANOVAs (repeated measures design) were performed, followed by Tukey’s multiple comparisons test.

**Supplemental Figure 4.**
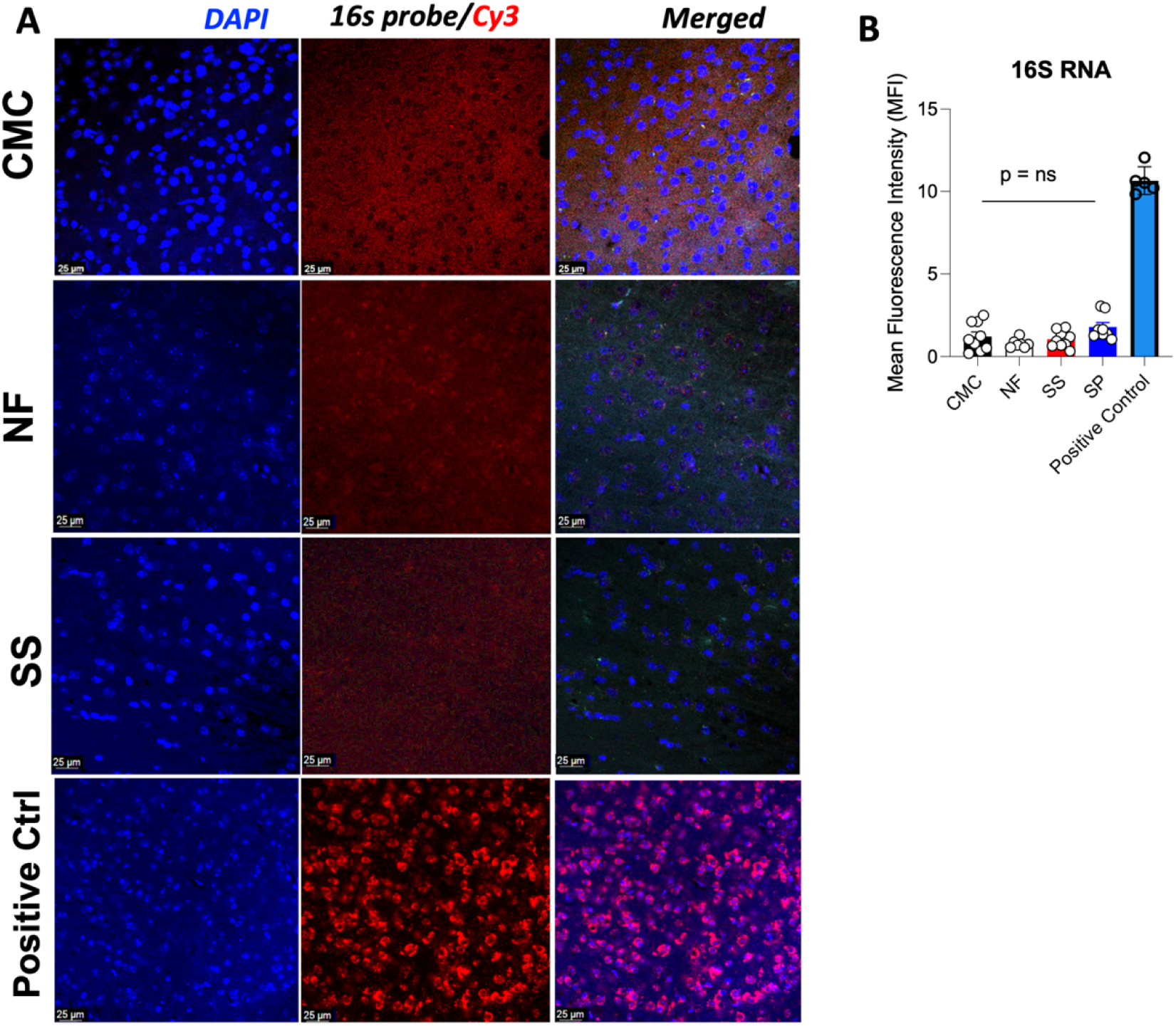
*S. salivarius* and *N. flavescens* nucleic acids were not detected in mouse brains by RNA-based in situ hybridization. **(A)** Representative images of sagittal mouse brain sections (5 μm FFPE) subjected to RNAScope in situ hybridization. Using a 16s probe, fluorescent labeling showed no detectable translocation of *S. salivarius* or *N. flavescens* RNAs from the oral cavity to the brain (red, Cy3 channel), with DAPI counterstain in blue. A positive probe (PPIB, a housekeeping gene) was used as a control (as shown in Figure 5 using brains from *S. parasanguinis*-treated mice). Images were acquired after screening the whole sagittal brain section at 40× magnification on a Zeiss LSM 510 Meta confocal microscope (1024 × 1024; n = 4). **(B)** Mean fluorescence intensity (MFI) was calculated using ImageJ/Fiji, and bars indicate mean ± SEM.

**Supplemental Figure 5.**
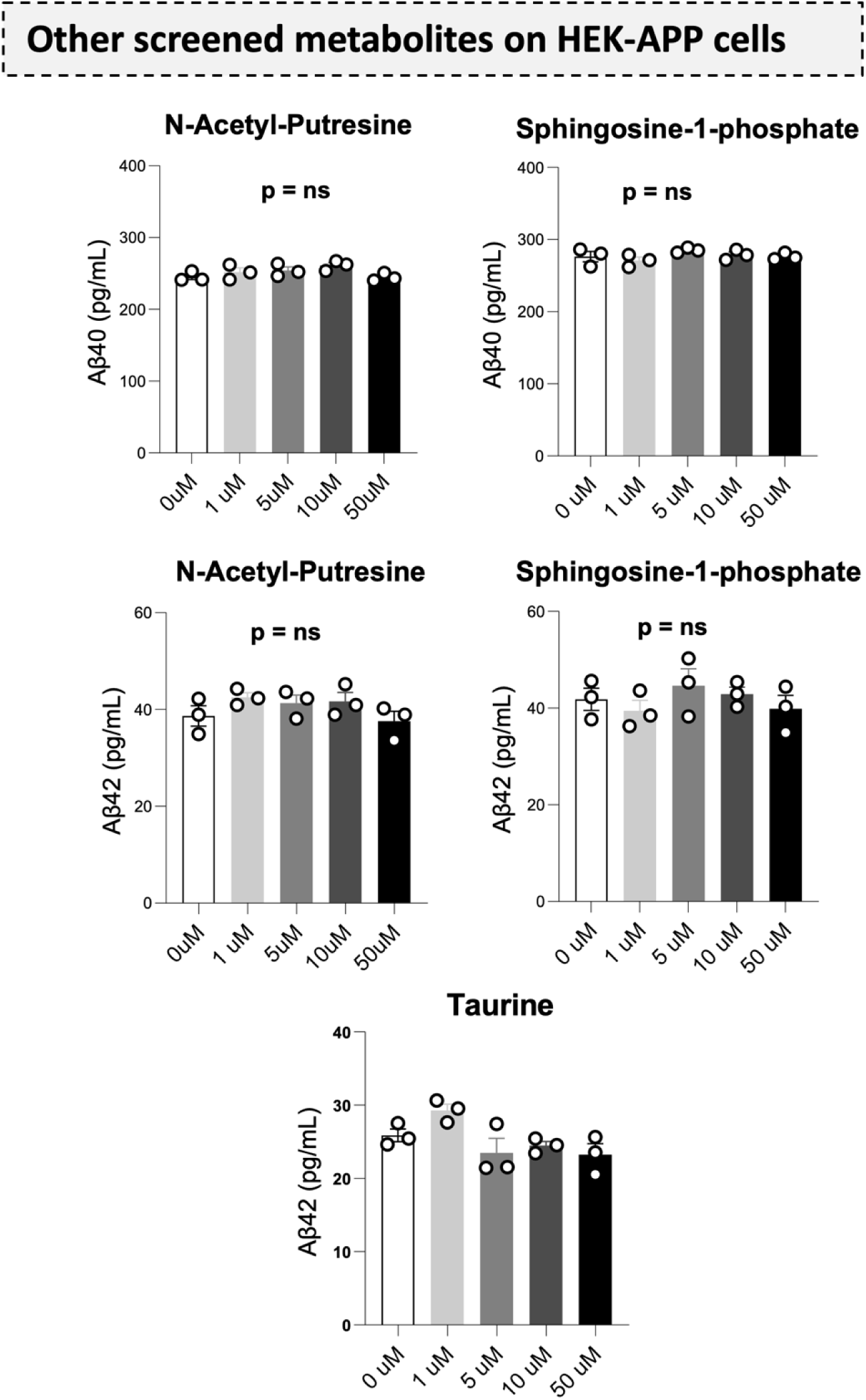
Screening of some shared metabolites detected in bacterial supernatant, mouse saliva, and brains on Aβ40 and Aβ42 levels. N-acetyl-putrescine, sphingosine-1-phosphate, and taurine were screened in HEK-APP cells at the indicated concentrations to monitor the Aβ40 and Aβ42 levels. None of them showed any prominent changes in the levels at any concentration. Data are presented as mean ± SEM (triplicate), following one-way ANOVA with Tukey’s post hoc test.

**Supplemental Figure 6.**
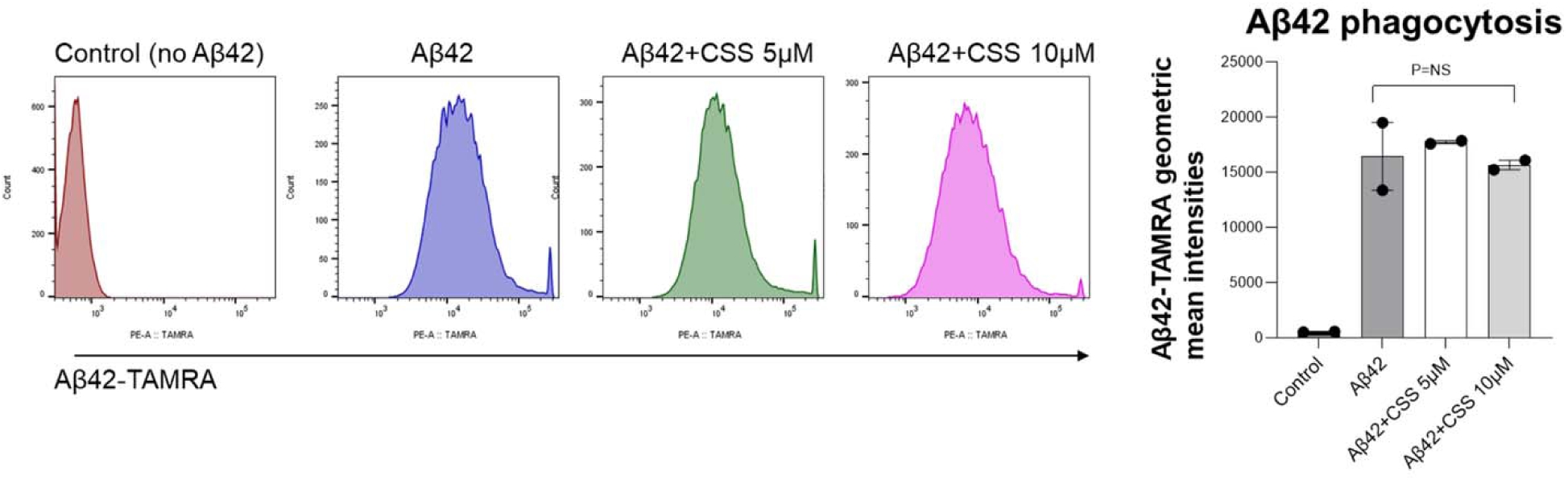
Cysteine s-sulfate treatment does not affect microglial phagocytosis of. **A**β**42.** Human iPSC-derived microglia were treated with or without cysteine s-sulfate for 24 hours in vitro. Aβ42 phagocytosis was analyzed by flow cytometry, as shown in one representative figure and a summarized figure.

**Supplemental Figure 7.**
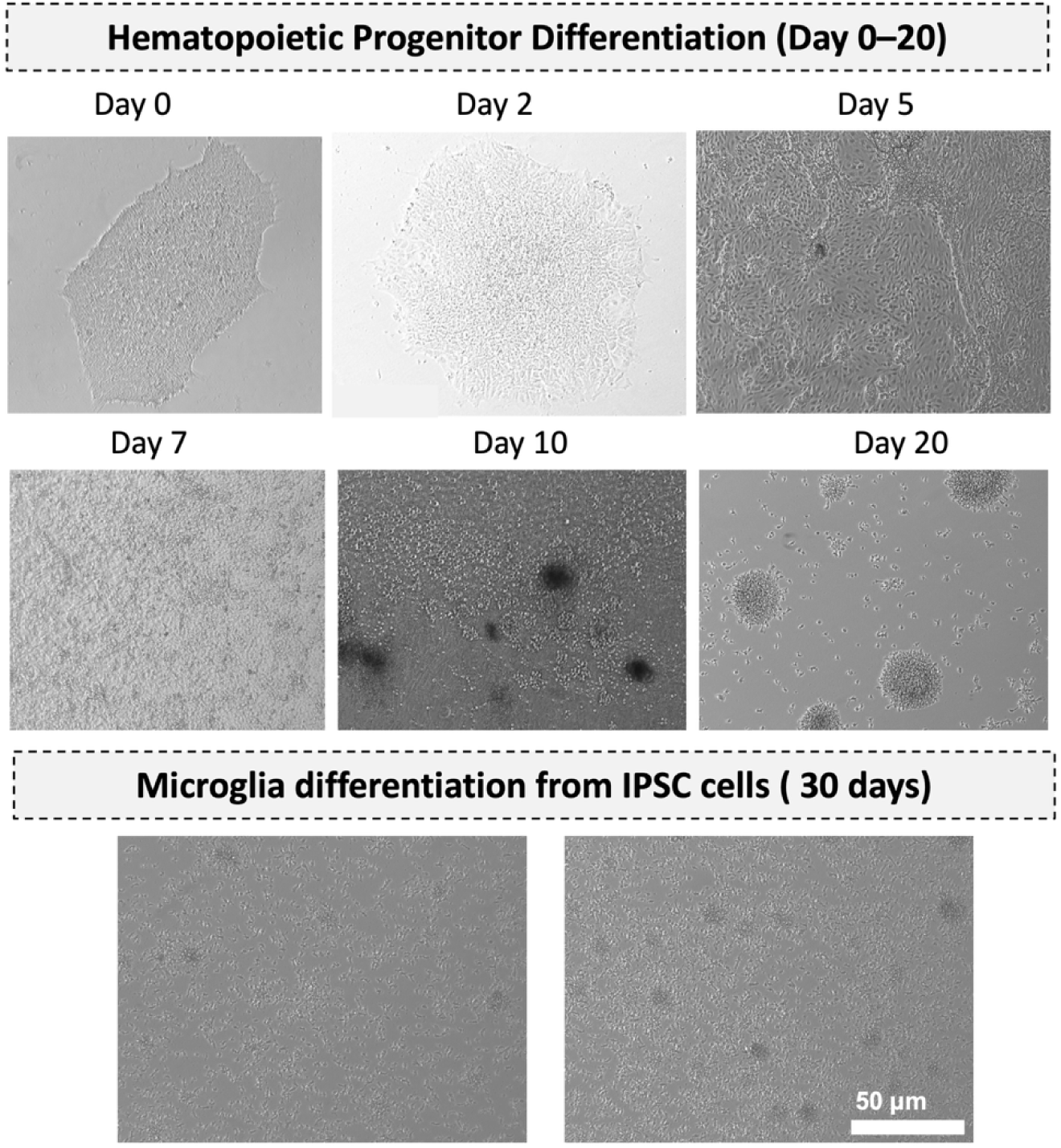
Images captured from Day 0 to Day 3 showing hematopoietic progenitor differentiation and microglial differentiation from the iPSCs.

